# Natural history of eukaryotic DNA viruses with double jelly-roll major capsid proteins

**DOI:** 10.1101/2024.03.18.585575

**Authors:** Mart Krupovic, Jens H. Kuhn, Matthias G. Fischer, Eugene V. Koonin

## Abstract

The phylum *Preplasmiviricota* (kingdom *Bamfordvirae*, realm *Varidnaviria*) is a broad assemblage of diverse viruses with comparatively short double-stranded DNA genomes (<50 kbp) that produce icosahedral capsids built from double jelly-roll major capsid proteins. Preplasmiviricots infect hosts from all cellular domains, testifying to their ancient origin and, in particular, are associated with six of the seven supergroups of eukaryotes. Preplasmiviricots comprise four major groups of viruses, namely, polintons, polinton-like viruses (PLVs), virophages, and adenovirids. We employed protein structure modeling and analysis to show that protein-primed DNA polymerases (pPolBs) of polintons, virophages, and cytoplasmic linear plasmids encompass an N-terminal domain homologous to the terminal proteins (TPs) of prokaryotic PRD1-like tectivirids and eukaryotic adenovirids that are involved in protein-primed replication initiation, followed by a viral ovarian tumor-like cysteine deubiquitinylase (vOTU) domain. The vOTU domain is likely responsible for the cleavage of the TP from the large pPolB polypeptide and is inactivated in adenovirids, in which TP is a separate protein. Many PLVs and transpovirons encode a distinct derivative of polinton-like pPolB that retains the TP, vOTU and pPolB polymerization palm domains but lacks the exonuclease domain and instead contains a supefamily 1 helicase domain. Analysis of the presence/absence and inactivation of the vOTU domains, and replacement of pPolB with other DNA polymerases in eukaryotic preplasmiviricots enabled us to outline a complete scenario for their origin and evolution.

**Significance:** Structural modeling of protein domains using advanced artificial intelligence-based methods such as AlphaFold2 may lead to insights into evolutionary relationships among proteins that are unreachable by sequence analysis. We applied this approach to elucidate the evolutionary relationships of four major groups of eukaryotic viruses: polintons, polinton-like viruses (PLVs), virophages, and adenovirids. We identified previously uncharacterized protein domains predicted to be essential for virus genome replication. Analysis of the presence/absence and inactivation of these domains suggests a complete scenario for the origin and evolution of this major part of the eukaryotic virosphere.

## Introduction

The phylum *Preplasmiviricota* (kingdom *Bamfordvirae*, realm *Varidnaviria*) is an expansive assemblage of viruses with moderate-sized double-stranded DNA (dsDNA) genomes (<50 kbp) that produce virions with icosahedral capsids (40–80 nm) assembled from double jelly-roll major capsid proteins (DJR-MCPs) (1). Preplasmiviricots infect hosts from all cellular domains, testifying to their ancient origin and evolutionary success. Indeed, the prokaryotic preplasmiviricots appear to predate the last universal cellular ancestor (LUCA) (2), whereas association with eukaryotic hosts has been mapped to the onset of eukaryogenesis (3). Extant preplasmiviricots have been detected in six of the seven supergroups of eukaryotes (3) and include four prominent groups of viruses currently referred to as polintons (*Mavericks*), polinton-like viruses (PLVs), virophages, and adenovirids (1).

Polintons (currently class *Polintoviricetes*) were first described as large, self-synthesizing transpos<u>ons</u> encoding a protein-primed family B DNA <u>pol</u>ymerase (pPolB) and a retrovirid-like <u>int</u>egrase (RVE-INT) as well as a cysteine protease of the Ulp1 deubiquitinylase family (vUlp1) and a genome packaging ATPase (Fig. 1) (4, 5). However, subsequent identification of the conserved genes for the major and minor capsid proteins (MCP and mCP, respectively) left little doubt that these elements are better described as endogenous viruses rather than transposons (6–9). Polintons are widespread across the eukaryotic evolutionary tree, being present in both diverse protists and animals, including vertebrates (7, 8, 10). Analysis of metagenomic databases using the polinton MCPs as queries led to the discovery of the distinct, highly diverse group of currently unclassified polinton-like viruses (PLVs) which, unlike bona fide polintons, mostly lack the pPolB and RVE-INT genes and instead encode other types of replication proteins and, occasionally, tyrosine superfamily integrases (11–13). However, a variant of PLVs, known as Tlr1 element (14), described in a ciliophore *(Tetrahymena thermophila* Nanney & McCoy, 1976), encodes RVE-INT but instead of pPolB encodes a large multidomain protein containing a C-terminal superfamily 1 helicase (S1H) domain (15), referred to as the ‘Tlr1-like helicase’. Other PLVs encode superfamily 3 helicases (S3H) fused to either DNA polymerase (DNAP) of family A (PolA) or archaeo-eukaryotic primase-polymerases (AEP) (11). At least two PLVs have been cultured, including Tetraselmis viridis virus S1 (TvV-S1) and Gezel-14T, both infecting marine algae (16, 17) (18). The third group of eukaryotic preplasmiviricots consists of virophages (currently class *Maveriviricetes*) have been discovered as hyperparasites of giant viruses assigned to bamfordviraen phylum *Nucleocytoviricota* (19–23). Virophages encode orthologous morphogenetic modules consisting of MCP and mCP, a packaging ATPase, and a vUlp1, but differ in terms of their genome replication proteins, with some virophages encoding pPolB (e.g., maviruses) and others encoding a PolA-S3H fusion (e.g., sputnikviruses) related to that found in some PLVs (24). Finally, the fourth group of eukaryotic preplasmiviricots consists of adenovirids (currently a distinct order in class *Tectiliviricetes*), a rather uniform group of viruses infecting diverse vertebrates, including humans (25). The gene content of adenovirids is similar to that of polintons, including pPolB, but they do not encode the typical genome packaging ATPase, which is replaced by a distinct ABC ATPase (26), nor RVE-INT and, indeed, do not integrate into host chromosomes.

**Figure 1.**
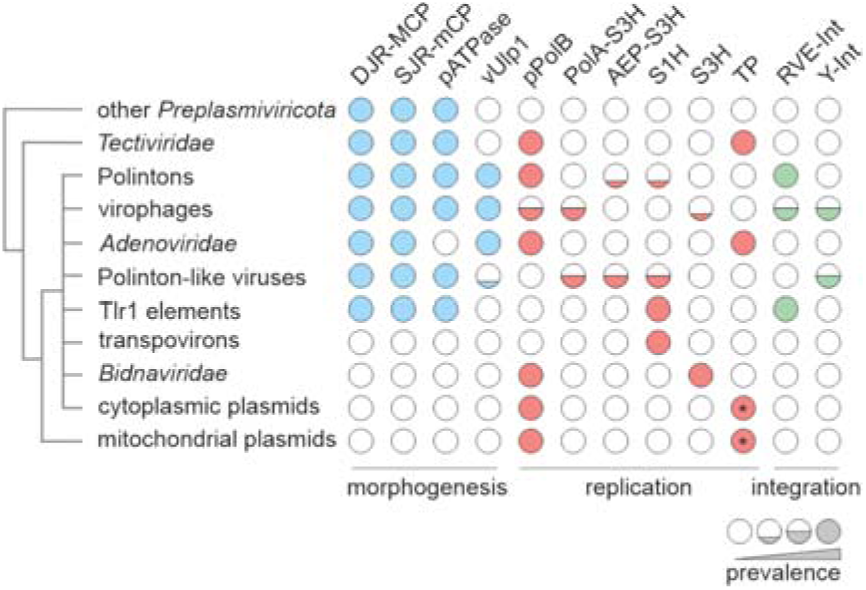
Relationships among viruses and non-viral mobile genetic elements encoding protein-primed family B DNAPs (pPolBs). The schematic tree on the left depicts relationships among the depicted elements and is loosely based on the pPolB phylogenies (9, 52). Circles depict gene presence and absence in the corresponding viruses and plasmids. AEP, archaeo-eukaryotic primase-polymerase; DJR-MCP, double jelly-roll major capsid protein; pATPase, FtsK-family genome packaging ATPase; PolA, family A DNAP; RVE-Int, retrovirid-like integrase; S1H, superfamily 1 helicase; S3H, superfamily 3 helicase; SJR mCP, single jelly-roll minor capsid protein; TP, terminal protein; Ulp1-PRO, Ulp1-family cysteine protease; Y-Int, tyrosine superfamily integrase. Asterisks indicate that the terminal proteins are encoded as part of pPolBs.

The genome replication proteins, i.e., pPolB and Tlr1-like helicase, connect preplasmiviricots to eukaryotic non-viral mobile genetic elements (Fig. 1) (27). In particular, two distinct groups of linear plasmids replicating, respectively, in the cytoplasm and mitochondria of fungi and in mitochondria of plants, encode pPolBs (9), whereas transpovirons, a group of linear plasmid-like molecules parasitizing mimivirids (28, 29), encode Tlr1-like helicases. Finally, the polinton pPolB gene has been horizontally transferred to insect parvovirids yielding a group of viruses with linear single-stranded DNA (ssDNA) genomes classified into family *Bidnaviridae* (30).

The evolutionary trajectory that led to the emergence and diversification of eukaryotic preplasmiviricots and their non-viral relatives remains unresolved. Phylogenetic analysis of the pPolB sequences suggested that bacterial viruses of the *Tectiviridae* family (currently *Preplasmiviricota*: *Tectiliviricetes*) gave rise to two lineages of eukaryotic elements, mitochondrial linear plasmids and polintons, respectively, with all other groups of eukaryotic viruses and plasmids evolving directly from polintons (9). Alternatively, it has been suggested that polintons evolved from mavirus-like virophages (20). Phylogenetic reconstructions based on the structural proteins yielded contrasting results, with polintons being at the base of a clade which included all eukaryotic viruses with DJR-MCPs in one analysis (31), but emerging as a terminal branch in another study, with virophages occupying the basal position (32). Given the fast evolution of capsid proteins and non-uniform selection pressures elicited by different hosts (e.g., exposure to humoral immunity), interpretation of the results of phylogenetic analyses based on structural proteins might not be straightforward.

Here, we explore evolution of eukaryotic preplasmiviricots and related non-viral elements through detailed structural analysis of their major replication proteins. Structural dissection showed that the pPolB of polintons carries a terminal protein domain implicated in protein-primed replication initiation, followed by a vOTU-like cysteine deubiquitinylase domain likely responsible for cleavage of the terminal protein from the rest of the pPolB polypeptide. Analysis of the presence/absence as well as inactivation of the protease domains in the homologs encoded by eukaryotic preplasmiviricots and related plasmids enabled us to establish the most likely sequence of events in their evolution.

## Results and discussion

### Complex domain architecture of eukaryotic preplasmiviricot pPolBs

The pPolBs encoded by preplasmiviricots vary widely in length, with the proteins of polintons being nearly twice as long as the pPolBs of tectivirids and mavirus-like virophages (Fig. 2A). To clarify the relationship between these pPolB homologs, we modeled and compared the pPolB structures from representatives of each virus group, namely, bacterial tectivirid Enterobacteria phage PRD1, Caenorhabditis briggsae polinton 1 (P1-CB), frog adenovirus 1 (FrAdV-1), mavirus virophage and Bombyx mori bidensovirus (BmBDV; *Bidnaviridae*), as well as those of mitochondrial and cytoplasmic linear plasmids (Fig. 2B). Analysis of the structural models showed that all the polymerases contain the characteristic TPR1 and TPR2 subdomains within their palm domain, confirming that they belong to the protein-primed subgroup of family B DNAPs. In the pPolB of the caudoviricete Bacillus subtilis phage phi29, the only pPolB for which the structure has been determined experimentally, TPR1 mediates interaction with the TP that primes genome replication, whereas TPR2 endows pPolB with the processivity and strand-displacement activities (33, 34). Notably, mavirus pPolB contained only a remnant of the exonuclease domain (Fig. 2B), which is conserved in all other family B DNAPs, suggesting that the mavirus polymerase has lower fidelity than other pPolBs. Unlike most other known pPolB-encoding viruses, mavirus encodes an S3H (Fig. 1), which likely unwinds DNA during genome replication.

**Figure 2.**
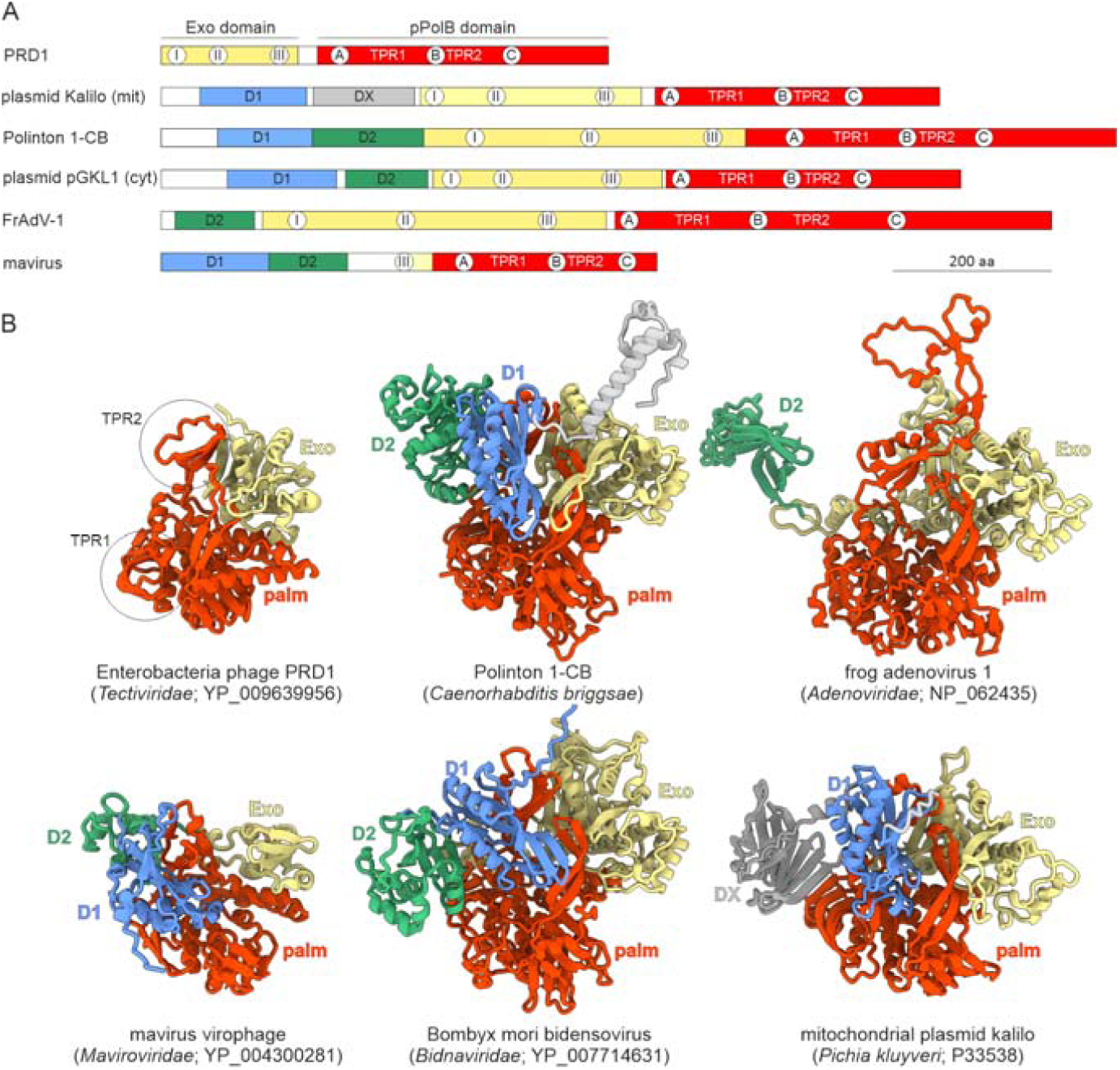
The domain organization of pPolBs encoded by preplasmiviricots and related elements. A. Schematic domain organization of pPolBs with homologous domains shown with matching colors. The locations of conserved motifs of the exonuclease and polymerization domains are indicated within the corresponding circles. The locations of the TPR1 and TPR2 subdomains are also shown. B. Structural models of pPolBs encoded by different groups of viruses and plasmids, with distinct domains colored using the same scheme as in panel A. The TPR1 and TPR2 subdomains characteristic of pPolBs are encircled. The models colored using pLDDT quality scores are shown in Figure S1, whereas the structural model of pPolB of yeast (*Kluyveromyces lactis*) plasmid pGKL1 is shown in Figure S2.

All eukaryotic pPolBs contain extended N-terminal regions which have no counterparts in pPolBs of prokaryotic viruses. Structural analysis of these regions revealed that they encompass two globular domains, tentatively labeled D1 and D2 in Figure 2B. D1 is absent in the adenovirid pPolB, whereas pPolBs of mitochondrial plasmids lack D2. Notably, the pPolB of yeast (*Pichia kluyveri*) plasmid kalilo instead of D2 contains an unrelated domain, DX, which is predicted to consist of ten β-strands with an alternating up-and-down orientation and has no significant similarity to proteins with known structures (Figure 2B). Despite lacking the exonuclease domain, the mavirus pPolB contained both D1 and D2, strongly suggesting that it is a derivative rather than the ancestor to the pPolBs of polintons and other elements.

### D1 domain is homologous to the terminal protein of PRD1-like tectivirids and adenovirids

As the pPolB name gives away, protein-primed polymerases work in tandem with TPs that are covalently linked through a phosphodiester bond to a nucleotide that functions as a primer for DNA synthesis. All three alcoholic amino acids (Ser, Thr, Tyr) were experimentally identified as linking or priming residues in TPs associated with different pPolBs (35, 36). Tectivirid and adenovirid TPs are encoded as separate proteins whereas in cytoplasmic linear plasmids, the TP is fused to pPolB and is proteolytically cleaved off following replication initiation by an unknown protease (37). The N-terminal regions of the polinton and bidnavirid pPolBs were similarly predicted to function as TPs (9, 30). TPs encoded by viruses from different families generally lack sequence similarity (36) and a high-resolution TP structure is only available for the caudoviricete Bacillus subtilis phage phi29 that is unrelated to preplasmiviricots (38). The phi29 TP has an elongated α-helical fold (Fig. S3) (38) and is clearly unrelated to the D1 and D2 domains of eukaryotic pPolBs that are predicted to adopt α/β folds with a central β-sheet (Fig. 2B, see below). To explore the structural relationships among the TPs of preplasmiviricots, we modeled the TP structures of tectivirids Enterobacteria phage PRD1 and Bacillus phage Bam35c, and adenovirid FrAdV-1.

Unexpectedly, the experimentally characterized TPs of PRD1 (39) and Bam35c (40) were predicted to have different folds. Although both proteins encompass a central β-sheet surrounded by α-helices, the topology of the β-sheet domains is different. Notably, the N-terminal elongated α-helical region of the Bam35c TP resembles the corresponding region of the phi29 TP (Fig. S3) although no overall structural similarity between these domains was observed. By contrast, the TPs of Enterobacteria phage PRD1 and FrAdV-1 are predicted to adopt the same fold, which is also found in the D1 domain conserved in the pPolBs of eukaryotic preplasmiviricots and plasmids (Fig. 3A). This fold consists of a 4-stranded β-sheet, which in some members is extended by an additional β-strand, and two α-helices (Fig. 3B). The TPs of both PRD1 and adenovirids contain N- and C-terminal α-helical extensions that are lacking in the D1 domains, suggesting that these regions mediate the interactions between the stand-alone TPs and the pPolB catalytic domain. In addition, adenovirid TPs contain an extensive α-helical insertion between β1 and α1, which is lacking in the PRD1 TP (Fig. 3A,B). The experimentally identified linking residues in the PRD1 (Tyr190) and adenovirid (Ser580) TPs occupy equivalent positions, between α2 and the C-terminal β4 strand, within non-structured loops. In all D1 domains, we identified either Ser or Tyr residues in positions equivalent to those of linking residues in the PRD1 and adenovirid TPs (Fig. 3C). Although the sequences surrounding the (predicted) linking residues were dissimilar, they all contained a helix-breaking Gly or Pro in the vicinity of the linking residue (Fig. 3D). By contrast, the negatively charged residues previously suggested to be a characteristic feature accompanying the linking residues based on the analysis of experimentally validated TPs (36) were not consistently present (Fig. 3D). Indeed, whereas the linking residue Ser580 in the human adenovirus (41) is flanked by Asp residues at the -2 and +2 positions, the homologous region in FrAdV-1 lacks both Asp residues, whereas Gly at the +1 position is conserved in both (Fig. 3D). Collectively, these observations suggest that D1 domains of eukaryotic pPolBs are homologous to the TPs of tectivirids and adenovirids and accordingly are responsible for the priming of DNA synthesis, solving a long-standing puzzle in the field.

**Figure 3.**
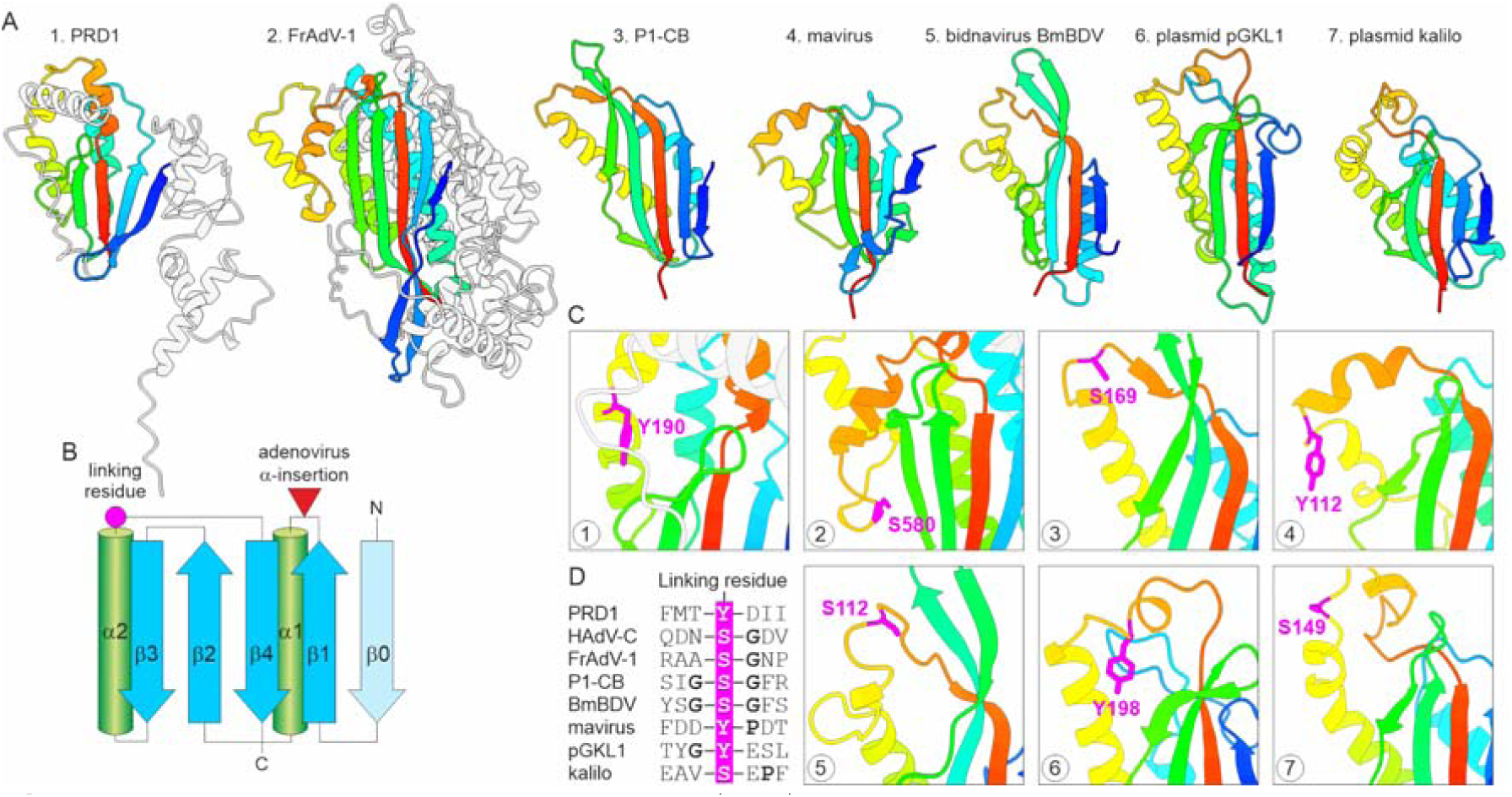
Conservation of terminal proteins (TPs). A. Structural comparison of the terminal proteins (TPs) of Enterobacteria phage PRD1 and frog adenovirus 1 (FrAdV-1) with D1s of pPolBs encoded by eukaryotic viruses and plasmids. Models are colored using the rainbow scheme from N-terminus (blue) to C-terminus (red). Non-conserved regions of the Enterobacteria phage PRD1 and FrAdV-1 TPs are shown in grey. B. Topology diagram of the PRD1-like TPs. The β-strand present in only some TPs is colored light blue and labeled β0. The positions of the linking residue and large α-helical insertion in adenovirids are indicated. C. Zoom-in on the (predicted) linking residues. The linking residues are shown in magenta using stick representation. Circled numbers in the corner of each box correspond to those in panel A, except for adenovirids (box 2), where a zoom-in on the experimentally characterized human adenovirus type C (HAdVC) TP is shown. D. Alignment of the regions flanking the predicted linking residues. Gly and Pro residues are shown in bold. Abbreviations: nematode (*Caenorhabditis briggsae* (Dougherty and Nigon, 1949)) polinton 1 (P1-CB); Bombyx mori bidensovirus (BmBDV).

### PRD1-like TPs likely evolved from the LF/PAD domain of family Y DNAPs

The lack of structural similarity between the PRD1, Bam35c, and phi29 TPs suggests that TPs have originated on at least three independent occasions. To gain insights into the provenance of the PRD1-like TPs, we performed searches queried with the structural models of TPs and pPolB D1 domains of preplasmiviricots and related elements against the PDB database using DALI. In all cases, the best hits with significant Z scores were to the little finger (LF) domain, known as the polymerase-associated domain (PAD) in eukaryotic proteins (Table S1). LF/PAD is a unique non-catalytic domain exclusively found at the C-terminus of family Y DNAPs, which are involved in mutagenic DNA repair in all domains of life (42–44). The LF/PAD domain is attached to the catalytic polymerase Y domain through an extended flexible linker and binds DNA across the major groove (42) (Fig. 4). It has been suggested that LF/PAD plays an important role in determining the enzymatic properties of the Y family DNAPs enzymes, modulating their processivity, ability to bypass template lesions, and capacity to generate base pair substitutions versus single-base deletions (45).

**Figure 4.**
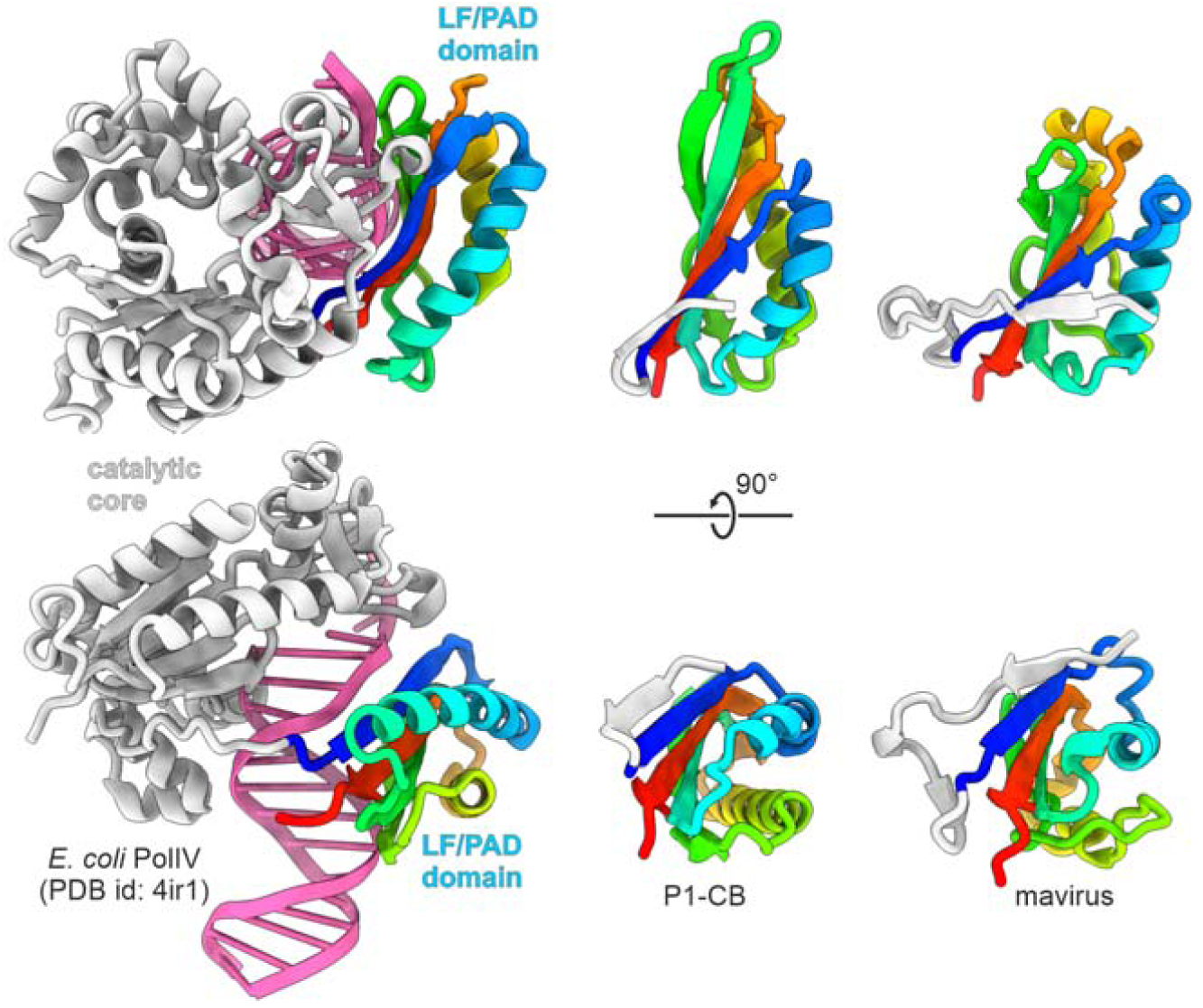
Origin of PRD1-like terminal proteins from the LF/PAD domain of family Y DNAPs. Structural comparison of the DNA-bound LF/PAD domain of the *Escherichia coli* DNAP IV (PolIV; PDB ID: 4IR1) with the TP domains of nematode (*Caenorhabditis briggsae* (Dougherty and Nigon, 1949)) polinton 1 (P1-CB) and mavirus virophage. The LF/PAD and TP domains are colored using the rainbow scheme from N-terminus (blue) to C-terminus (red), whereas the catalytic domain of PolIV and dsDNA are shown in grey and magenta, respectively.

Comparison of the PRD1-like TP models with the structures of the LF/PAD (Fig. 4) confirmed that the two groups of domains have the same fold consisting of a 4-stranded β-sheet and two α-helices (Fig. 3B). The conservation of the domain organization of the Y-family DNAPs across all domains of life testifies to the ancient association between the catalytic domain of Y-family DNAPs and LF/PAD, likely traceable to LUCA. Conversely, the sporadic distribution of LF/PAD-like TPs in bacterial viruses likely reflects a more recent recruitment. Thus, we hypothesize that the PRD1-like TP evolved from LF/PAD by gaining the ability to covalently bind the priming nucleotide, a facile adaptation given the simplicity of the sequence and structural contexts of the linking residue, which itself can vary (Fig. 3D). Notably, similar to Y family DNAPs, the TP domain is linked to the catalytic palm domain of pPolB in eukaryotic preplasmiviricots through an extended unstructured linker, which likely enables flexibility and movement. Conceivably, the preexisting capacity of the LF/PAD to interact with damaged DNA facilitated its recruitment as a terminal protein to initiate replication of linear DNA genomes.

### D2 domain is a vOTU-like cysteine protease

The fusion of the TP and catalytic pPolB domains raises the question whether and how the two domains are uncoupled during processive DNA synthesis, following the initiation of DNA replication. In the case of cytoplasmic linear plasmids, such as pGKL1 of *Kluyveromyces lactis* (yeast), it has been demonstrated that the 26 kDa N-terminal priming domain is proteolytically cleaved off by an unknown protease and stays covalently linked to the 5’ of the nascent DNA strand (46). By contrast, in certain mitochondrial linear plasmids, such as the kalilo plasmid of *Pichia kluyveri* (yeast), the mass of the terminal protein (120 kDa) corresponds to the entire DNAP, suggesting that the priming domain is not detached from the catalytic palm domain (47).

DALI searches queried with the structural models of the pPolB D2 domains from eukaryotic preplasmiviricots and related elements (Fig. 5A) yielded significant hits to diverse viral and cellular ovarian tumor domain (OTU) deubiquitinylases (Table S1, Fig. 5B), a superfamily of cysteine proteases with a distinct variant of the papain-like fold, also known as otubains (Fig. 5C) (48, 49). The viral OTU (vOTU) homologs process viral polyproteins and also function as deubiquitinylases counteracting antiviral defenses (50). We hypothesize that the vOTU domains of pPolBs are responsible for the proteolytic detachment of the TP domains from the catalytic domains.

**Figure 5.**
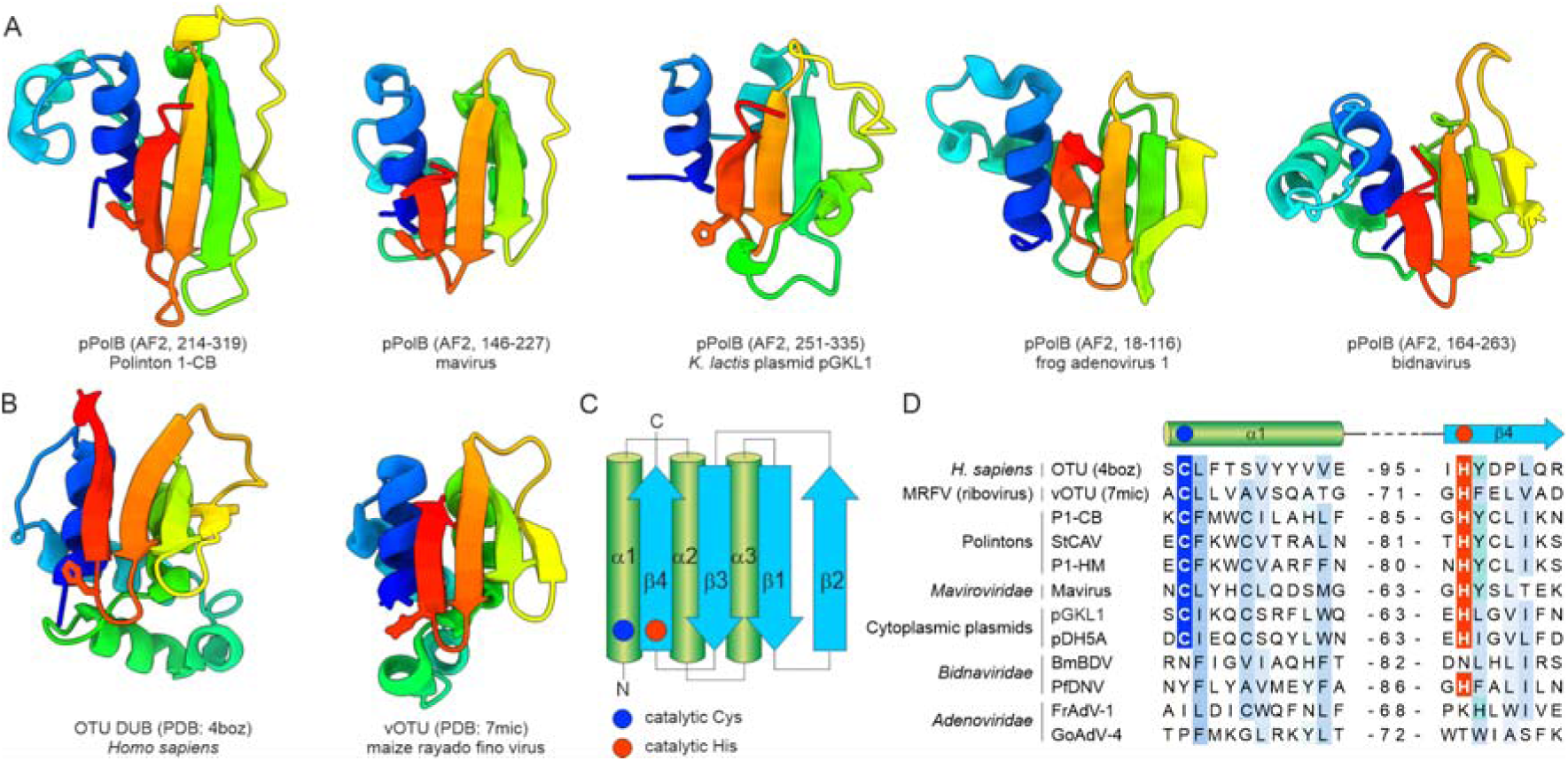
Conservation of the vOTU-like cysteine protease domain in pPolBs. A. Structural comparison of the vOTU domains present in pPolBs. B. Crystal structures of OTU from humans (PDB: 4BOZ) and vOTU from an RNA virus, maize rayado fino virus (PDB: 7MIC). Models are colored using the rainbow scheme from N-terminus (blue) to C-terminus (red). When present, the catalytic Cys and His residues are shown using stick representation. C. Topology diagram of the (v)OTU domains, with the catalytic Cys and His residues shown as blue and red circles, respectively. D. Sequence alignment of the α1 and β4 elements harboring the catalytic Cys (blue background) and His (red background) residues, respectively. The numbers in the middle denote the spacing between the two structural elements. BmBDV, Bombyx mori bidensovirus; FrAdV-1, frog adenovirus 1; GoAdV-4, goose adenovirus 4; P1-CB, nematode (*Caenorhabditis briggsae* (Dougherty and Nigon, 1949)) polinton 1; P1-HM, hydra (*Hydra vulgaris* Pallas, 1766) polinton 1; pDH5A, yeats (*Debaryomyces hansenii*) plasmid pDH5A; pGKL1, yeast (*Kluyveromyces lactis*) plasmid pGKL1; StCAV, Stylophora coral adintovirus; PfDNV, Phylloscopus fuscatus densovirus.

The invariant catalytic Cys and His residues located within the α1 and β4 elements of the vOTUs, respectively (Fig. 5C), are conserved in polintons, mavirus virophage and cytoplasmic plasmids (Fig. 5D), strongly suggesting that these are active proteases. To remove the TP domain from the palm domain, vOTU could cleave either immediately downstream of the TP domain or downstream of the vOTU itself. The experimentally determined size of the TP domain of the pPolB of *Kluyveromyces lactis* plasmid pGKL1 (46) is consistent only with the former scenario, i.e., cleavage between the TP and vOTU domains.

Although the pPolBs of adenovirids and bidnavirids contain vOTU-like domains, they encompass replacement of the predicted active site residues. Conceivably, the split of the TP-encoding fragment into a stand-alone TP gene in adenovirids rendered the protease activity of vOTU domain obsolete, subsequently leading to its inactivation. By contrast, bidnavirids appear to employ a strategy akin to that of mitochondrial linear plasmids, whereby the entire pPolB remains covalently attached to the nascent DNA strand. Indeed, analysis of the structural proteins of bidnavirid particles showed the presence of a 118-kDa protein corresponding to pPolB (51), consistent with the inactivation of the vOTU domain in bidnavirids.

### Tlr1-like helicases are multidomain proteins containing pPolB domains

We have previously observed that Tlr1-like helicases encoded by PLVs and transpovirons (Fig. 1) contain sequence motifs similar to the catalytic motifs of pPolBs (Fig. 6A) (52). The availability of powerful structure-prediction tools, such as AlphaFold2 (53), now enables scrutinizing this observation in greater detail and further exploring the domain organization of these large proteins. We modeled the structures of the Tlr1-like helicases of Tlr1 element (15) and transpoviron Mama associated with mimivirid Acanthamoeba castellanii mamavirus (28) (Fig. 6B, Fig. S1C). The models confirmed that Tlr1-like helicases, in addition to S1H domains, contain palm domains with the characteristic DNAP catalytic motifs (Fig. 6C). Comparison of the palm domains of RNA-primed and protein-primed PolBs with the palm domains of Tlr1-like helicases unequivocally shows that the latter belong to the protein-primed variety and contain the diagnostic TPR1 and TPR2 subdomains (Fig. 6D). Previously, we suggested that changing the first of the two Asp residues in the polymerase C motif (DTD) involved in metal coordination would render the palm domain inactive (52). However, the same Asp residue is missing in the PolBs of mavirus virophage (Fig. 6C), mitochondrial circular plasmids (54), pipolins (55), and certain archaeal DNAPs (56), all of which are nevertheless catalytically active. By contrast, motifs A and B are conserved. Thus, we predict that Tlr1-like helicases possess DNAP activity and are functionally analogous to other polymerase-helicase fusion proteins, such as PolA-S3H and AEP-S3H.

**Figure 6.**
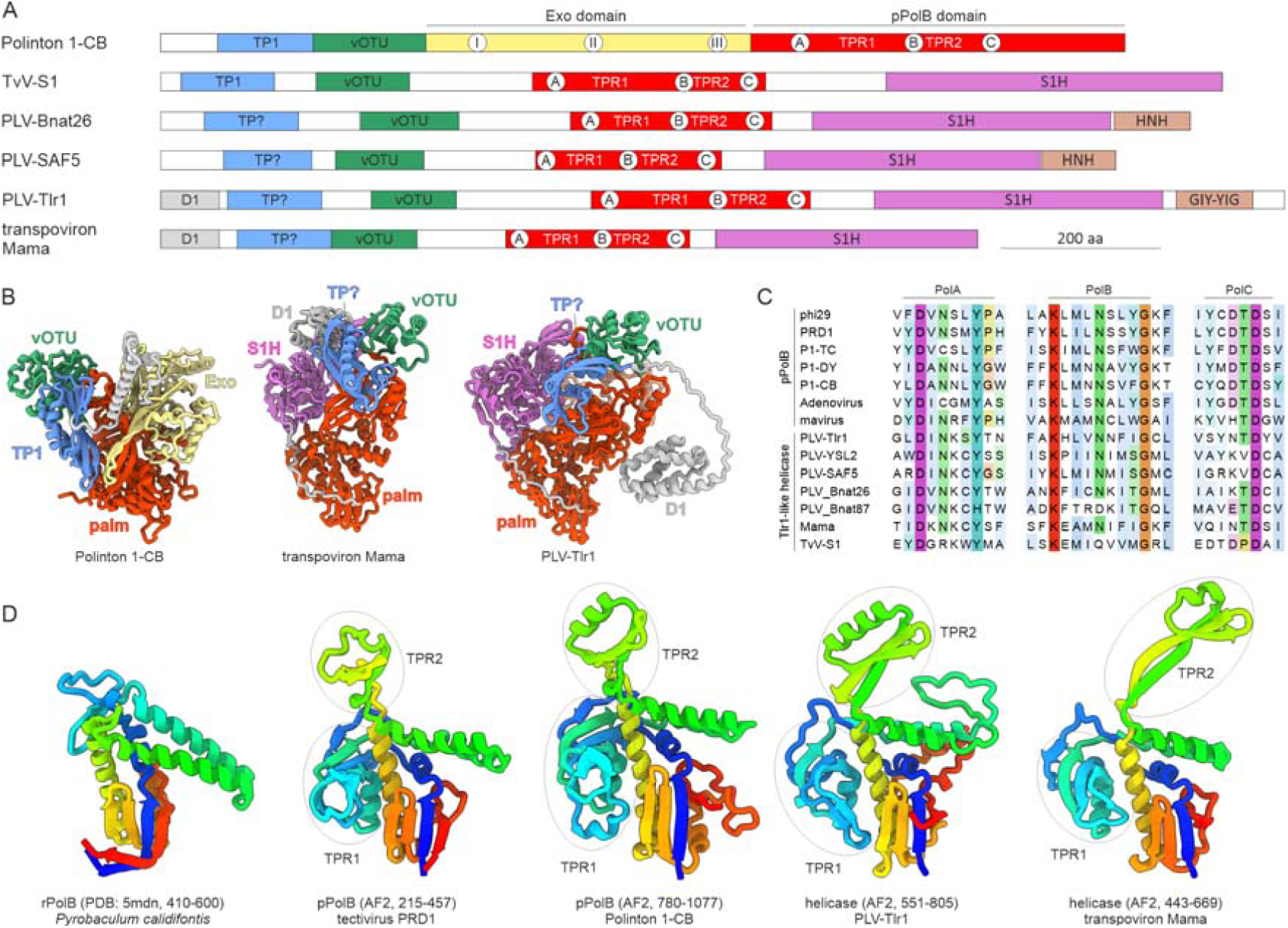
The domain organization of Tlr1-like helicases. A. Schematic domain organization of Tlr1-like helicases encoded by PLVs and transpovirons with homologous domains shown with matching colors. The annotated domain organization of the pPolB of nematode (*Caenorhabditis briggsae* (Dougherty and Nigon, 1949)) polinton 1 (P1-CB) is shown for comparison. The locations of conserved motifs of the exonuclease and polymerization domains are indicated within the corresponding circles. The locations of the TPR1 and TPR2 subdomains are also shown. Question marks denote uncertainty. B. Structural models of Tlr1-like helicases encoded by transpoviron Mama and Tlr1 element, with distinct domains colored using the same scheme as in panel A. The model of pPolB of polinton 1-CB is shown for comparison. The models colored using the pLDDT quality scores are shown in Figure S1. C. Sequence alignment of the motifs, PolA–C, conserved in family B DNAPs. D. Structural comparison of the palm domains of RNA-primed and protein-primed family B DNAPs (rPolBs and pPolBs, respectively) with those of Tlr1-like helicases. Models are colored using the rainbow scheme from N-terminus (blue) to C-terminus (red). The TPR1 and TPR2 subdomains characteristic of protein-primed polymerases are circled. The coordinates of the depicted regions are indicated in parentheses.

Phylogenetic analysis of the palm domain of Tlr1-like helicases suggested that it evolved from polinton pPolBs (52). Similar to the latter, Tlr1-like helicases contain extended N-terminal regions (Fig. 6A) that thus far evaded functional annotation. Analysis of the structural models of the PLV-Tlr1 and transpoviron Mama helicases showed that both contain three globular domains upstream of the palm domain. The N-terminal domains are highly variable in different PLV and transpoviron helicases and none are similar to proteins with known structures in the PDB database. By contrast, the domains proximal to the palm domains were identified as vOTUs (Fig. S4A), with both catalytic Cys and His residues conserved in all analyzed PLV and transpoviron sequences (Fig. S4B), suggesting that these vOTUs are active proteases, likely involved in proteolytic processing of the Tlr1-like helicases, as proposed above for the pPolBs of eukaryotic preplasmiviricots. Unexpectedly, the topology of the middle domain, occupying a position equivalent to that of the TP in pPolBs was variable, with at least three different variants among PLVs, although all were predicted to adopt α/β folds, with central β-sheets (Fig. S5). In particular, the corresponding domain in Tlr1 was unique, whereas the one in transpoviron Mama and some PLVs have a distinct fold resembling the Bam35c TP, although convergence in this case cannot be excluded. Notably, TvV-S1 contains a bona fide PRD1-like TP domain, suggesting that in this virus the middle domain functions as TP. Given that in S1H phylogenies, TvV-S1 occupied a basal position with respect to other PLVs and transpovirons (52), the PRD1-like TP domain is the likely ancestral state in the evolution of PLVs that was subsequently replaced or permutated into structurally distinct but functionally equivalent domains. Finally, at the C-terminus, Tlr1-like helicases of different PLVs contain homing endonucleases of two distinct families, GIY-YIG and HNH (Fig. 6A). Notably, polinton pPolBs also contain diverse homing endonuclease domains inserted at different positions of the protein (10), suggesting that these insertions in pPolB and Tlr1-like helicases result from intein invasions.

### Evolutionary history of *Preplasmivirocota*

Dissection of the pPolB and Tlr1-like helicase structures described here prompted a scenario for the evolution of preplasmiviricots and related non-viral elements (Fig. 7). The presence of structurally distinct TPs in tectivirids and their uniformity in most of the eukaryotic pPolBs suggest that eukaryotic preplasmiviricots evolved from a particular subgroup of tectivirids with PRD1-like TPs. The fusion of the TP domain to the polymerase domain in all eukaryotic pPolBs except for adenovirids (see below), along with the association between preplasmiviricots and eukaryotes being traceable to the last eukaryotic common ancestors (LECA) (3), suggest that the fusion occurred early during eukaryogenesis or possibly in a yet undiscovered group of bacterial tectivirids that includes direct ancestors of eukaryotic preplasmiviricots. Phylogenetic analyses of eukaryotic pPolBs consistently suggest that the first split occurred between mitochondrial linear plasmids and the ancestor of the remaining eukaryotic pPolBs (Fig. 7) (9) and this conclusion is supported by our structural comparisons. Conceivably, this split coincided with the escape of the ancestor of polintons from the proto-mitochondrial bacterial endosymbiont into the cytoplasm of the emerging proto-eukaryote. In some mitochondrial linear plasmids (e.g., in rapeseed (*Brassica napus* L.) (57)), the TP domain is fused directly to pPolB, whereas in others (e.g., the kalilo plasmid of *Pichia kluyveri*) the two domains are separated by an additional domain (DX) of unknown function. The vOTU domain was recruited subsequently in only one of the two major branches of preplasmiviricots, the polintons, yielding a multidomain pPolB with the TP-vOTU-EXO-PALM domain organization that is also shared by cytoplasmic linear plasmids. Thus, polintons appear to have been the first group of eukaryotic preplasmiviricots to evolve from tectivirids, subsequently giving rise to the entire diversity of the eukaryotic preplasmiviricots. Phylogenetic analyses have confidently shown that cytoplasmic linear plasmids are derived from (proto)polintons through the loss of the morphogenetic module, switching to a non-viral mode of propagation.

**Figure 7.**
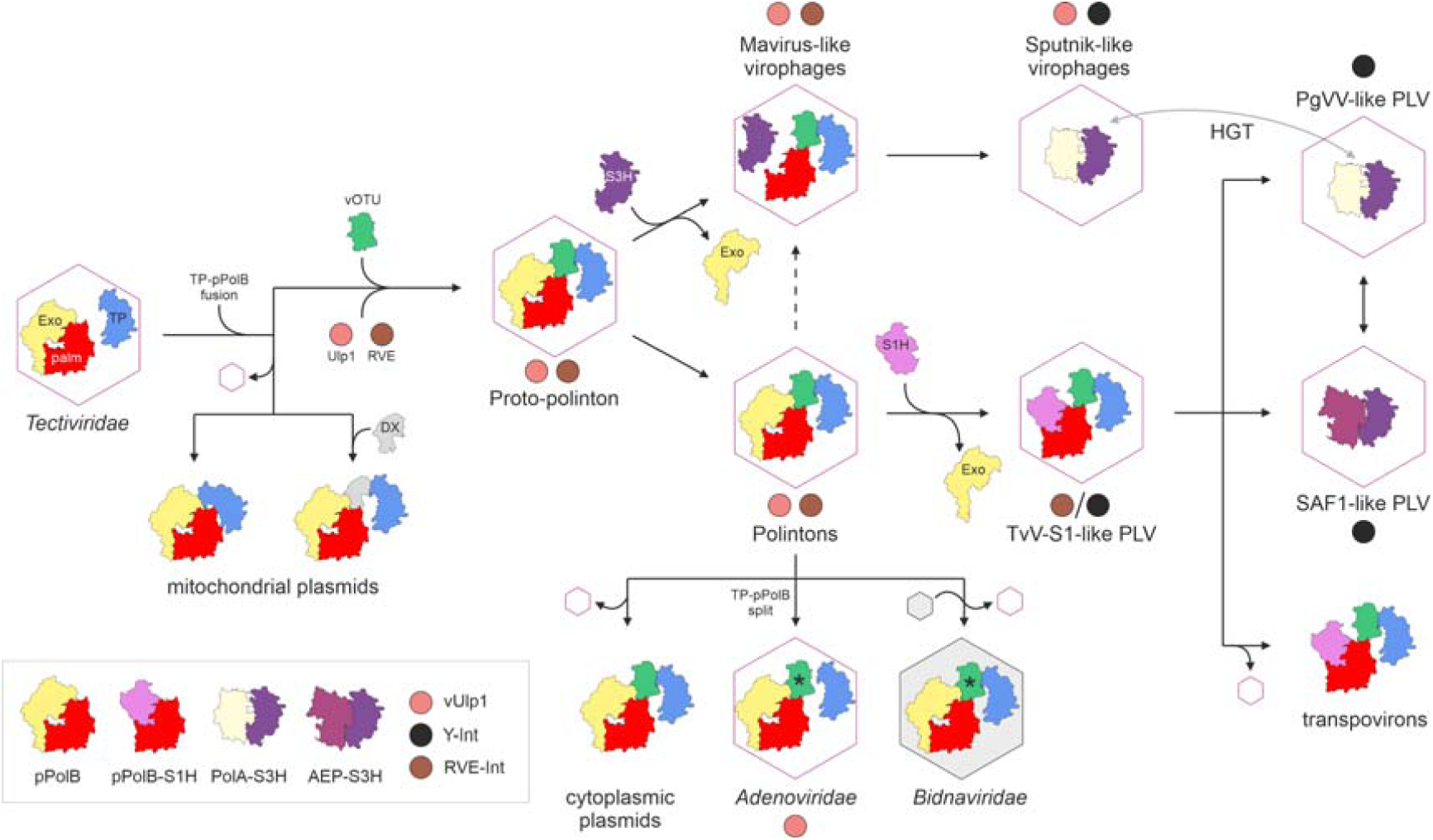
Natural history of preplasmiviricots and derived eukaryotic elements. The schematic depicts a proposed scenario for the evolution of eukaryotic preplasmiviricots and related elements from a bacterial tectivirid ancestor. Hexagons represent icosahedral capsids built from double jelly-roll major capsid proteins (DJR-MCPs). The grey hexagon indicates replacement of the DJR-MCP with an unrelated capsid protein in bidnavirids. Black and grey arrows indicate evolution and horizontal gene transfer (HGT), respectively. The dashed arrow denotes an alternative route of evolution. Asterisks indicate inactivation of vOTU domains. AEP, archaeo-eukaryotic primase-polymerase; Exo, exonuclease; PLV, polinton-like virus; PolA, family A DNAPs; pPolB, protein-primed family B DNAP; RVE-Int, retrovirid-like integrase; S1H, superfamily 1 helicase; S3H, superfamily 3 helicase; TP, terminal protein; TvV-S1, Tetraselmis viridis virus S1; (v)Ulp1, (viral) Ulp1-family cysteine protease; Y-Int, tyrosine superfamily integrase.

Adenovirids encode a PRD1-like TP as a stand-alone protein, resembling the organization observed in bacterial viruses. However, the presence of inactivated vOTU domains at the N-terminus of their pPolBs leaves no doubt that adenovirids evolved from polintons rather than inherited the split TP organization from bacterial viruses. The inactivated vOTU domain in adenovirids could have been repurposed for the interaction between the TP and pPolB or other replication factors. Notably, the split between the TP and vOTU domains seems to have occurred independently in different groups of polintons. For example, we identified a polinton in hood corals (*Stylophora pistillata* Esper, 1797) (8) (Fig. S6) in which the TP is encoded by a separate gene, whereas the catalytically active vOTU domain is fused to pPolB, as in adenovirids (Fig. 5D). Thus, adenovirids might have evolved from a group of polintons in which the TP was already detached from pPolB. The vOTU domain was also inactivated in bidnavirids which, however, retained the TP domain fused to the pPolB. Consistently, pPolB has been identified in the virions of bidnavirids (51), suggesting that, like in the case of some mitochondrial linear plasmids that lack the vOTU domain altogether, the pPolB is not processed and stays attached to the genomic DNA.

Polintons are also at the root of mavirus-like virophages and PLVs. The mavirus pPolB has a unique TP-vOTU-PALM domain organization, indicating that it is a highly derived DNAP that lost most of the exonuclease domain, ruling out the possibility that the mavirus pPolB is ancestral to that encoded by polintons. Nevertheless, given the high divergence of the virophage MCPs, it appears likely that virophages evolved from a proto-polinton ancestor although direct evolution from polintons cannot be ruled out either (Fig. 7). The mavirus-like pPolB resembles the ancestor of the Tlr1-like helicases widespread in PLVs, which have the TP-vOTU-PALM-S1H domain organization. We predict that the palm domains of these proteins are active DNAPs and accordingly, they should be more accurately denoted as pPolB-S1H fusion proteins, akin to PolA-S3H and AEP-S3H widespread in other viruses, rather than Tlr1-like helicases.

Notably, the loss of the exonuclease domain is accompanied by the gain of a helicase, S3H in maviruses (encoded by a separate gene; Fig. S6) and S1H fused to pPolB in PLVs. The two other polymerase-helicase fusions common in eukaryotic preplasmiviricots, AEP-S3H and PolA-S3H, also lack the exonuclease domain. Conversely, PolA encoded by diverse phages (e.g., T7) lack the helicase domain but contain the exonuclease domain (58). Thus, there seems to be a compensatory relationship between the exonuclease and helicase activities such that both increase the polymerase fidelity via different mechanisms. The presence of distinct helicases, S3H and S1H in mavirus and PLVs, respectively, and the pronounced divergence between the corresponding MCPs suggest that the exonuclease domains have been independently lost in the two virus groups. These findings clarify the position of PLVs and virophages in the evolution of *Preplasmiviricota*, suggesting that both virus groups diverged from a polinton-like ancestor with a bona fide pPolB.

The loss of the structural module yielded transpovirons, all of which encode pPolB-S1H replication proteins and thus apparently evolved from PLVs. The link between virophages, PLVs and transpovirons also extends to their mode of propagation. All known virophages and transpovirons and at least some PLVs, such as Phaeocystis globosa virus virophage (PgVV) (17, 59), are parasites of giant viruses of the phylum *Nucleocytoviricota*, suggesting that this interaction emerged early in the evolution of eukaryotic viruses. As noted above, besides the pPolB-S1H, some virophages and PLVs encode PolA-S3H or AEP-S3H replication proteins, respectively. In phylogenetic analyses based on the structural module, mavirus-like virophages are consistently basal to virophages encoding PolA-S3H (31, 32), suggesting that pPolB is the ancestral replication protein which was replaced in some virophage lineages. Indeed, the unique PolA-S3H fusion protein first discovered in sputnikviruses (24) has now been identified also in diverse PLVs (12), suggesting extensive exchange of the replication and structural modules between PLVs and virophages.

Taken together, these results clarify the evolutionary history of the *Preplasmiviricota* and reassert the central position of polintons in the evolution of eukaryotic viruses, plasmids, and transpovirons. The revised scenario outlined here calls for refinement of the *Preplasmiviricota* taxonomy.

## Materials and methods

### Protein structure prediction and analysis

Structural models of pPolBs encoded by mitochondrial and cytoplasmic plasmids were downloaded from the AlphaFold database (60). The structural model of the pPolB of polinton 1 from a nematode (*Caenorhabditis briggsae* (Dougherty and Nigon, 1949)) was taken from a previous study (10). The remaining protein structures were modeled using AlphaFold2 (53) or DeepFold (61) through ColabFold v1.5.5 (62) “alphafold2_multimer_v3” model with six recycles. To improve the quality of the model, structural templates were provided when available. Structures were searched against the PDB database using DALI (63). Comparison of predicted and experimentally resolved structures from PDB was performed using MatchMaker (64). Protein structures and structural models were visualized using ChimeraX v1.7.1 (65).

## Data availability

All protein structural models generated in this study are available in SI data file 1.

## Authors contributions

M.K. and E.V.K. designed research; M.K. performed research; M.K., M.G.F., J.H.K., and E.V.K analyzed data; M.K. and E.V.K. wrote the manuscript that was edited and approved by all authors.

## Acknowledgements and funding sources

This work was supported by a grant from l’Agence Nationale de la Recherche (ANR-23-CE02-0022) to M.K. E.V.K. is supported through the Intramural Research Program of the National Institutes of Health (National Library of Medicine). This work was supported in part through a Laulima Government Solutions, LLC, prime contract with the U.S. National Institute of Allergy and Infectious Diseases (NIAID) under Contract No. HHSN272201800013C. J.H.K. performed this work as an employee of Tunnell Government Services (TGS), a subcontractor of Laulima Government Solutions, LLC, under Contract No. HHSN272201800013C. The views and conclusions contained in this document are those of the authors and should not be interpreted as necessarily representing the official policies, either expressed or implied, of the U.S. Department of Health and Human Services or of the institutions and companies affiliated with the authors.

## SI figures and legends

**Figure S1.**
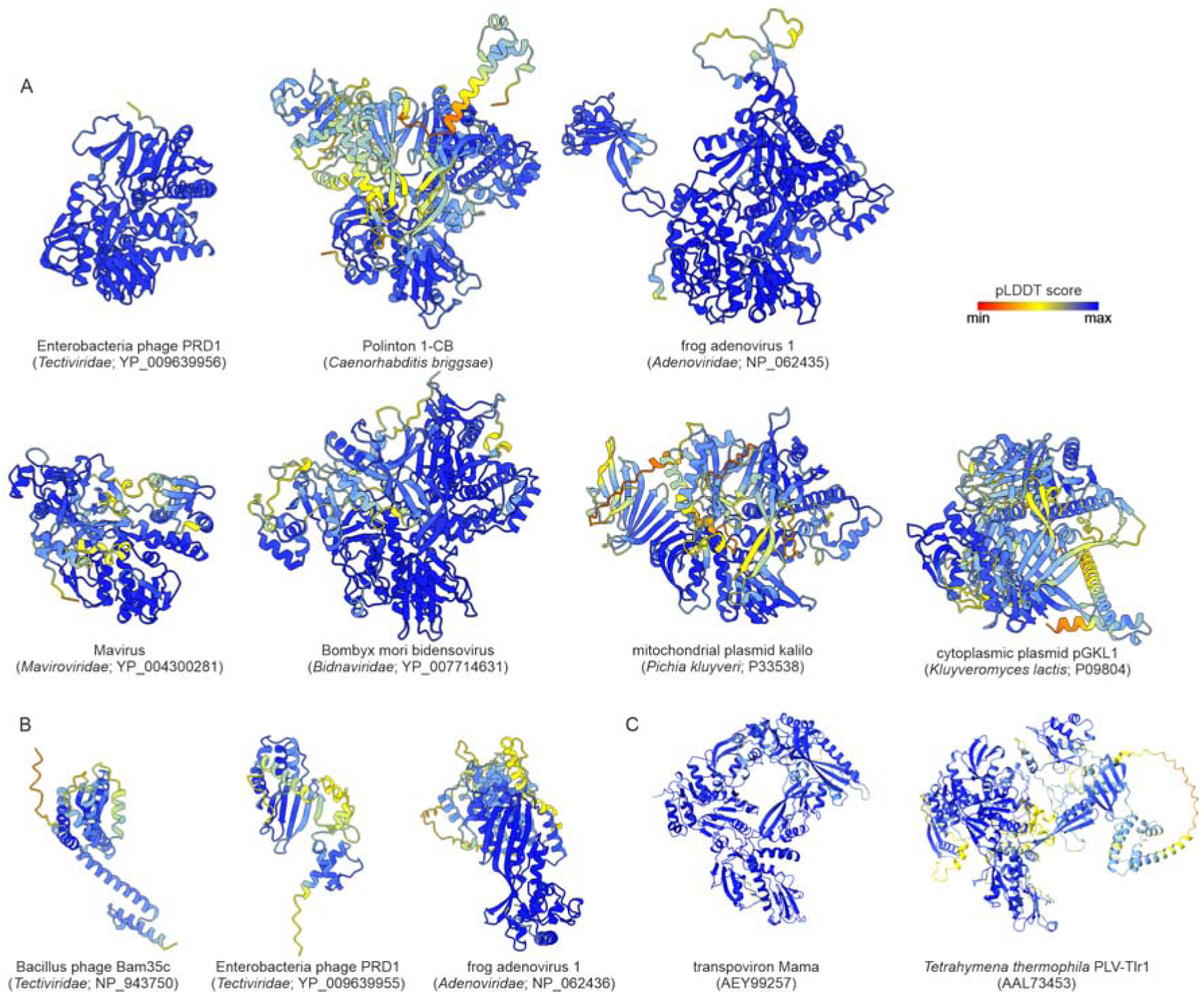
Structural models colored according to their per-residue confidence scores assessed by predicted local distance difference test (pLDDT). The pLDDT scale is show in in the top right corner of the figure. GenBank or UniProt accession numbers of the modeled proteins are provided in parentheses. A. Protein-primed family B DNAPs. B. Terminal proteins. C. Tlr1-like helicases.

**Figure S2.**
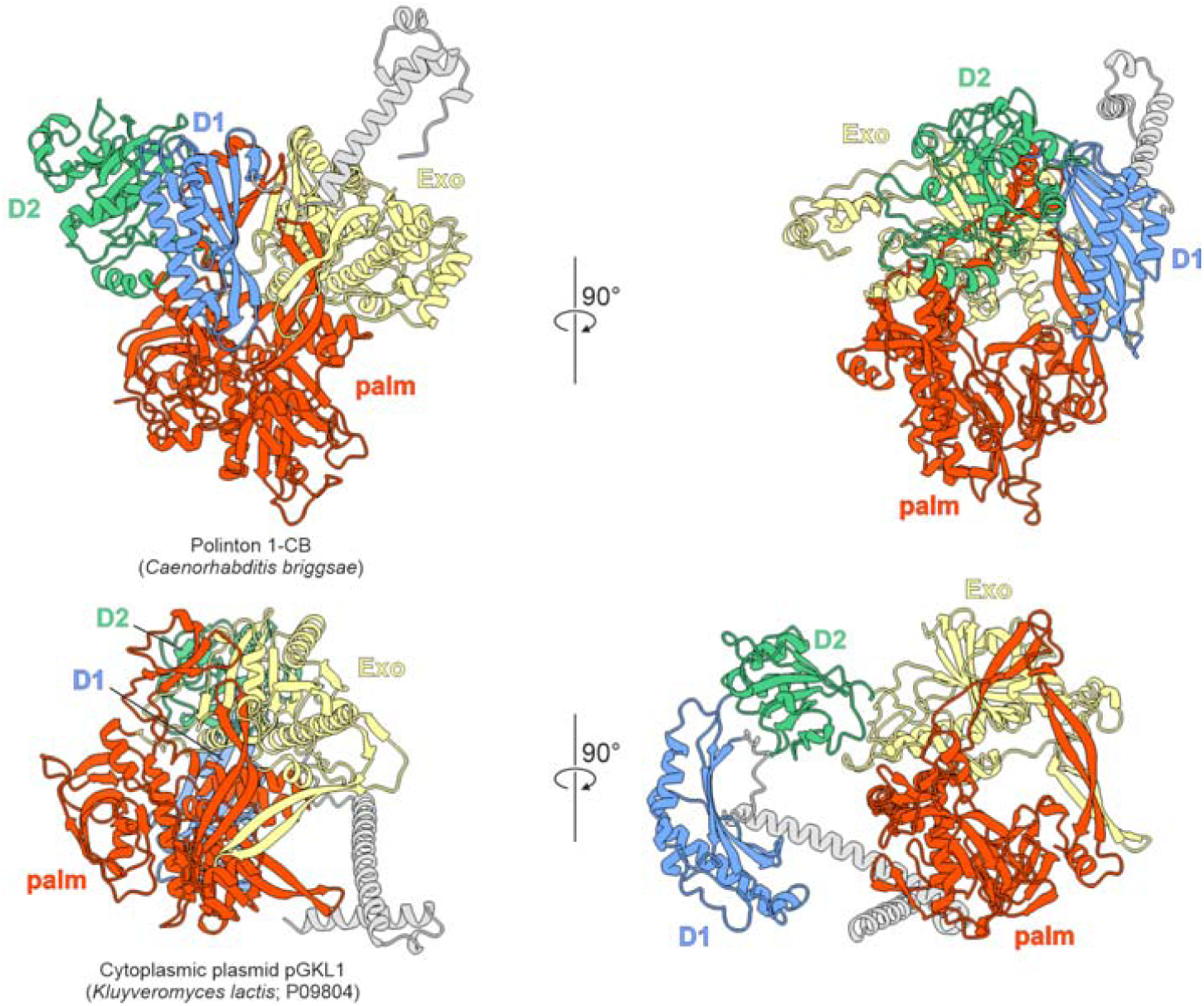
Comparison between structural models of the pPolBs encoded by polinton 1-CB and cytoplasmic linear plasmid pGKL1 from a yeast (*Kluyveromyces lactis*). The view on the left is in the same orientation as in Figure 2B, whereas the view on the right is rotated by 90°, so that D1 and D2 of pGKL1 pPolB become visible. Domains are colored as in Figure 2B.

**Figure S3.**
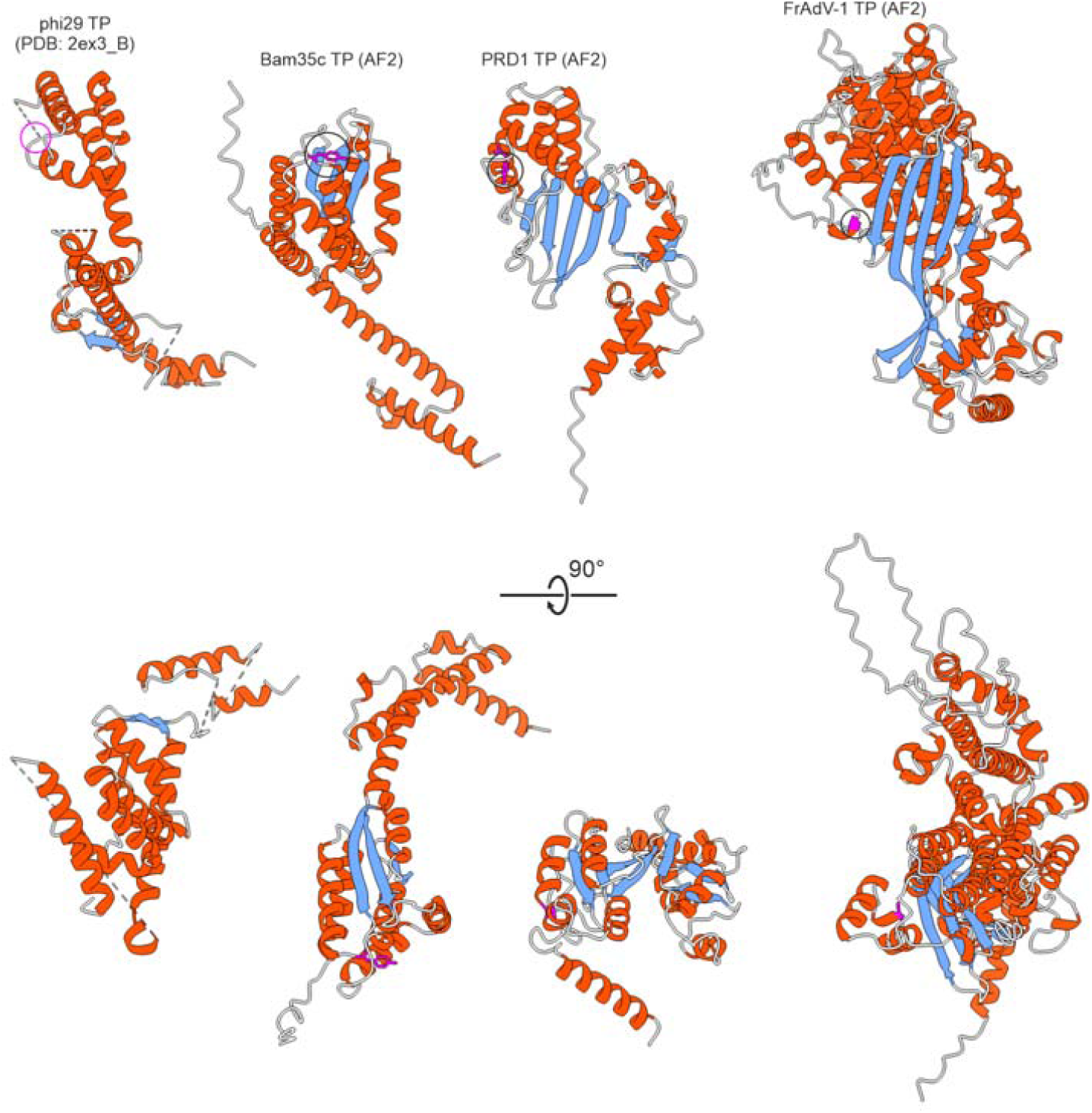
Structural comparison of terminal proteins. Comparison of the X-ray structure of the terminal protein (TP) of Bacillus subtilis phage phi29 with the structural models of terminal proteins encoded by tectivirids Bacillus phage Bam35c and Enterobacteria phage PRD1, and frog adenovirus 1. The structures are colored according to secondary structure elements: α-helices, red; β-strands, blue; coils, grey. The linking residues are circled. Note that the linking residue has not been resolved in the Bacillus subtilis phage phi29 structure (PDB: 2ex3_B).

**Figure S4.**
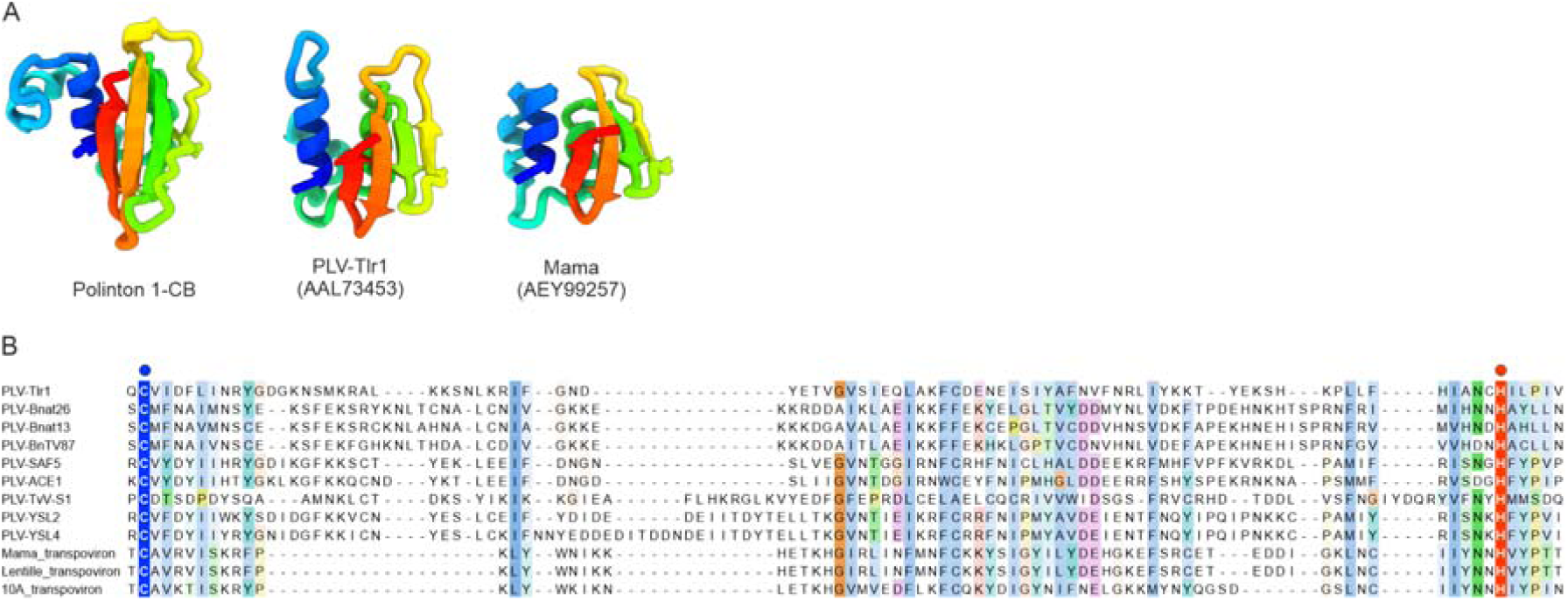
vOTU domains of Tlr1-like helicases. A. Comparison of the vOTU structural models from the pPolB of polinton 1-CB with the corresponding domains from the Tlr1-like helicases of PLV-Tlr1 and transpoviron Mama. The models are colored using the rainbow scheme from N-terminus (blue) to C-terminus (red). B. Sequence alignment of the vOTU domains of Tlr1-like helicases encoded by PLVs and transpovirons. The catalytic Cys and His residues are indicated with blue and red circles, respectively.

**Figure S5.**
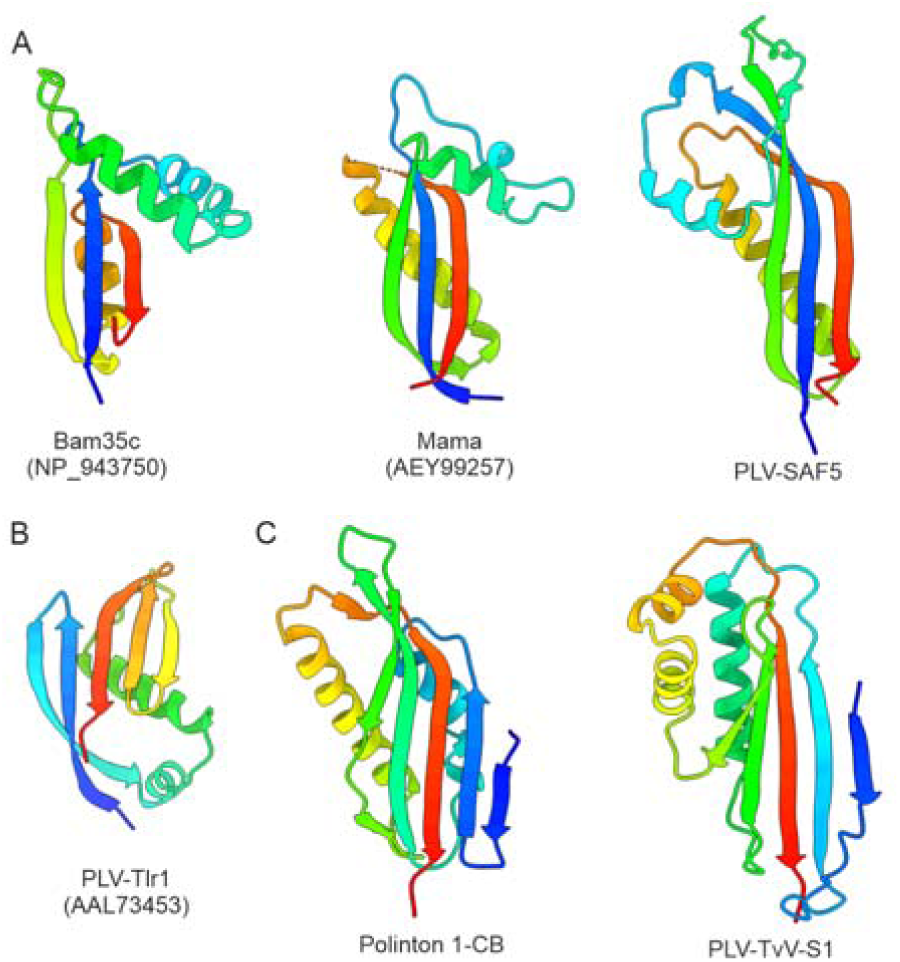
Structural models of the domains preceding the vOTU domain in Tlr1-like helicases and their comparison to the terminal proteins. Three structurally distinct domains are shown. A. Domain showing similarity to the terminal protein of tectivirid Bacillus phage Bam35c. B. Tlr1 element-specific domain. C. The domain of PLV TvV-S1 is compared to the PRD1-like terminal protein domain of polinton 1-CB. The models are colored using the rainbow scheme from N-terminus (blue) to C-terminus (red). PLV, polinton-like virus; TvV-S1, Tetraselmis viridis virus S1,

**Figure S6.**
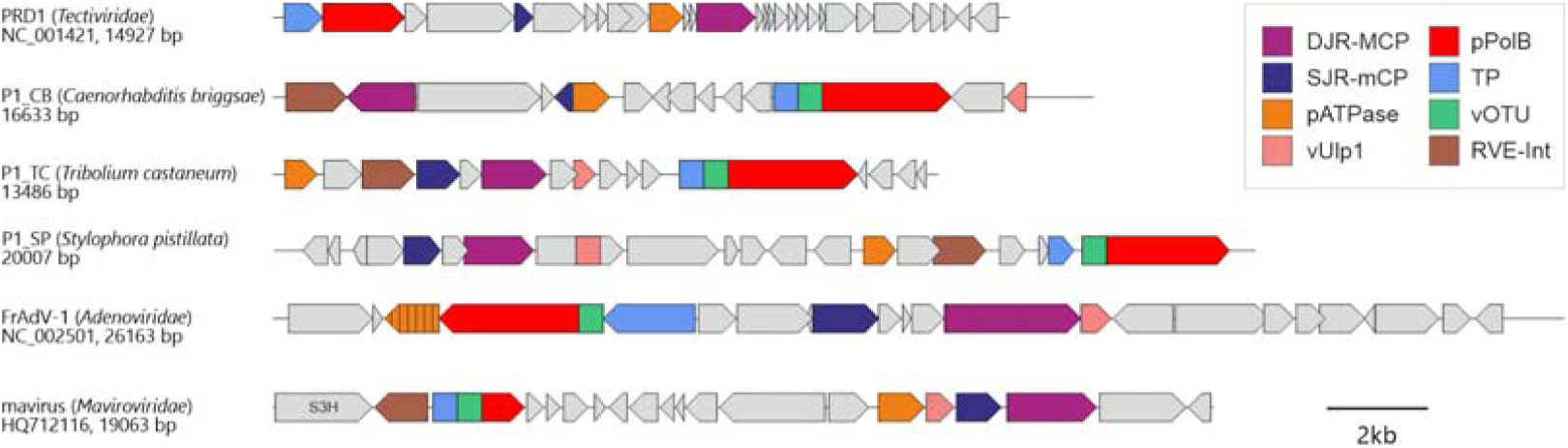
Genome maps of selected preplasmiviricots. Relevant genes are color-coded.

## SI table

**SI Table S1.**
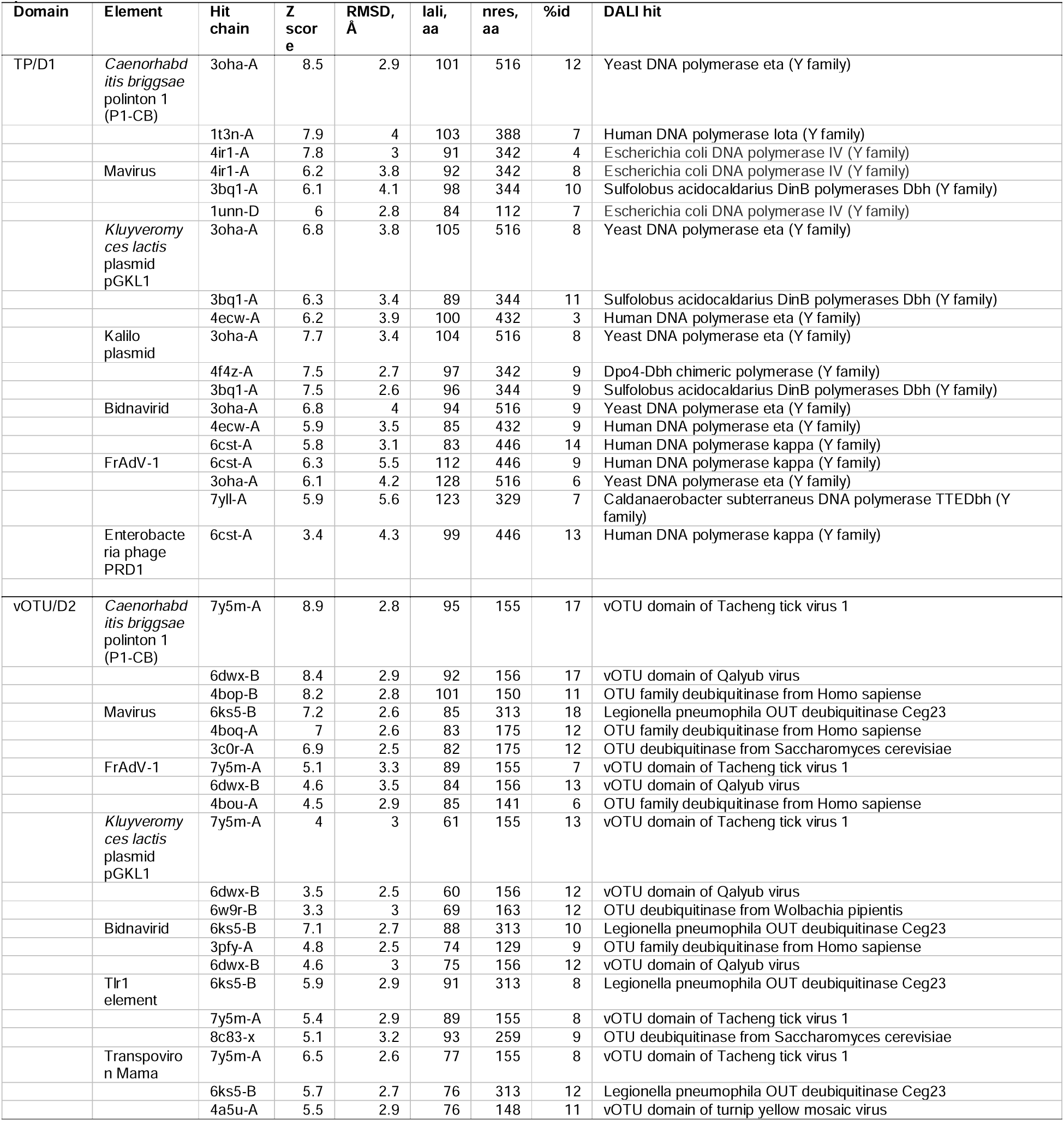
Results of the DALI searches queried with the structural models of the terminal protein (TP) and vOTU domains.

